# Physical activity in children and adolescents with Charcot-Marie-Tooth disease: A cross-sectional case-controlled study

**DOI:** 10.1101/493908

**Authors:** Rachel A. Kennedy, Kate Carroll, Kade L. Paterson, Monique M. Ryan, Joshua Burns, Kristy Rose, Jennifer L. McGinley

## Abstract

**Background:** Disability related to the progressive and degenerative neuropathies known as Charcot-Marie-Tooth disease (CMT) affects gait and function, increasing with age and influencing physical activity in adults with CMT. The relationship between disease, ambulatory function and physical activity in children and adolescents with CMT is unknown.

**Method:** A cross-sectional case-controlled study of 50 children with CMT and age- and gender-matched typically developing (TD) controls [mean age 12.5 (SD 3.9) years]. A 7-day recall questionnaire assessed physical activity; disease severity and gait-related function were measured to explore factors associated with physical activity.

**Results:** Children with CMT were less active than TD controls (estimated weekly moderate to vigorous physical activity CMT 283.6 (SD 211.6) mins, TD 318.0 (SD 202.5) mins; *p* < 0.001). The children with CMT had moderate disability [CMT Pediatric Scale mean score 20 (SD 8) /44] and reduced ambulatory capacity in a six-minute walk test [CMT 485.1 (SD 160.9) metres, TD 639.8 (83.1) metres; *p* < 0.001]. Physical activity correlated with greater disease severity (ρ = -0.52, *p* < 0.001) and six-minute walk distance (ρ = 0.71, *p* < 0.001).

**Conclusions:** Disease-related disability affects physical activity and gait-related function in children and adolescents with CMT compared to TD peers. Reduced physical activity adversely affects function across the timespan of childhood and adolescence into adulthood.

## Introduction

Physical activities involving walking, running and jumping are often limited by progressive muscle weakness caused by the inherited peripheral neuropathies collectively known as Charcot-Marie-Tooth disease (CMT) (1, 2). Reduced physical activity is associated with negative health outcomes irrespective of disability, with a compounding effect in adulthood (3–5). Children with physical disabilities are less physically active than their typically developing peers, with the potential for poorer health outcomes including obesity related to inactivity (6, 7). Likewise, for children and adolescents (“children”) with CMT, the negative health consequences of reduced physical activity are likely to compound the impairments of their disease. However, to date there have been no studies of physical activity in children with CMT.

The natural history of CMT is of progressive weakness and worsening of disease-related disability. Adults with CMT are less physically active than unaffected adults (8). Furthermore, adults with greater disease severity report being less active than adults with milder disease (9). Whilst adults with CMT are more likely to experience greater disease-related limitations to physical activity than children with CMT, weakness in CMT is evident from early childhood (10) and disease severity increases throughout childhood and adolescence (11). Therefore, it is possible that children with CMT are less active than their typically developing peers due to their disease and at risk of negative health outcomes related to inactivity.

Understanding physical activity in children’s daily lives requires a measurement tool that reflects activity throughout the day. There is no single method of measuring physical activity in children that captures both objective and descriptive information about the type of activity undertaken (12). A review of physical activity monitoring outlined several paediatric-specific self-report physical activity tools that can be completed within a single study visit and have been validated with objective measures of activity (accelerometers and activity monitors) (13). Self-report recall questionnaires record activities undertaken at school and out of school hours, during organised extra-curricular and week-end activities, and incidental activities such as active mobility in the community (13).

Disease-related impairments in children with CMT likely affect engagement in recreational and organised sports, as well as incidental activities such as using stairs and walking longer distances. Gait-related activities undertaken safely and sufficiently in the everyday environments of daily life can be defined as functional ambulation (14). It is likely that children with CMT are less physically active, and also report and demonstrate limitations to functional ambulation. Therefore, the aims of this study were to compare self-reported physical activity of children and adolescents with CMT to that of age- and gender-matched typically developing (TD) peers; and secondly, to investigate associations between physical activity, disease severity and measures of functional ambulation in children and adolescents with CMT.

## Methods

### Study design

A cross-sectional, case-controlled study was conducted across two specialist paediatric neuromuscular centres. Ethical approval was gained from The Royal Children’s Hospital (RCH) (HREC 33272 and 36019), The Sydney Children’s Hospitals Network (LNR/16/SCHN/132) and The University of Melbourne (1647763). Parents/guardians and participants 12 years and older were provided with a plain language information statement, and participant assent and parent/guardian written consent were gained prior to enrolment.

### Participants

Fifty ambulant children with a confirmed genetic or clinical diagnosis of CMT were recruited from either the Neuromuscular Clinic at The Royal Children’s Hospital (RCH), Melbourne, or the Peripheral Neuropathy Management Clinic at Children’s Hospital, Westmead (CHW), Sydney between January 2016 and October 2017. Fifty age- and gender-matched typically developing (TD) children were recruited from Melbourne and regional Victoria. Inclusion criteria included the ability to walk > 75 m without gait aids (orthotics permitted). Exclusion criteria included other developmental/ neuromuscular/ musculoskeletal disorders that could affect gait, and lower limb surgery or injury in the preceding 6 months that might limit physical activity or gait.

### Procedures and assessments

A single clinical assessment was conducted in the respective outpatient clinic. Participant descriptors and characteristics including date of birth, standing height and weight, dominant hand and foot, and CMT subtype (CMT only) were recorded and body mass index (BMI) calculated (kg/m^2^). Disease severity in the participants with CMT was assessed with the Charcot-Marie-Tooth disease Pediatric Scale (CMTPedS) an 11-item, well-validated and reliable, linearly-weighted outcome measure of disease and disability in children and adolescents aged 3-20 years which measures motor function, strength, and balance (15). The TD participants were assessed on items from the CMTPedS relating to lower limb function; specifically, dorsiflexor and plantar flexor strength, standing balance, standing long jump and the six-minute walk test (6MWT). Additional foot and ankle characteristics assessed in all participants included foot posture index (FPI) (16) and maximum dorsiflexion range in a weight bearing lunge (17).

### Physical activity questionnaire – child (PAQ-C)

The PAQ-C is a valid and reliable self-reported, 7-day recall measure of physical activity in children (18–20). The PAQ-C was used for all participants enrolled in this study irrespective of age, as it aligns with the typical daily activity schedule within the Australian school system (19). Participants were asked to report the types and intensity of physical activity in the seven days prior to their study visit. If the visit was undertaken during school holidays, the child was asked to reflect on the last seven days that they were at school, as extra-curricular activities are often suspended over holiday periods, altering physical activity patterns. Children aged 8 years and younger completed the PAQ-C with assistance of a parent or adult.

The PAQ-C consists of nine questions scored on a 5-point Likert scale (S1 Appendix) (19). The first relates to a checklist of physical activities, both sporting and leisure, common to children and adolescents. The participant was asked to rate how often they performed these activities in the past seven days on a 5-point scale from “none” to “7 or more”. Minor modifications were made to the checklist for this study to reflect activities common to Australia, in keeping with prior reports from other cultures (21–23). For example, activities related to snow sports were removed and Australian sports such as cricket and netball were added (see S1 Appendix for full list of activities). The remaining eight questions utilised a 5-point ordinal scale to assess the level of physical activity (none through to vigorous); during the school day (PE/sports class, recess and lunch breaks), immediately after school, in the evenings and on the week-end; a general statement relating to how physically active the child was during the past 7 days; and a general statement relating to physical activity levels for each day of the previous week.

The PAQ-C is scored out of five, with 1/5 reflecting low activity and 5/5 reflecting high activity (24). From the PAQ-C score, a group estimate of moderate to vigorous physical activity (MVPA) was derived from a method calibrated against accelerometry (25). This method was used to calculate a group estimate of weekly MVPA for the CMT and TD groups.

### Six-minute walk test

The 6MWT was utilised as a test of ambulatory capacity and was conducted according to the local clinic’s usual practice and following the CMTPedS protocol (15). In Melbourne, participants wore their own well-fitting footwear (typically a pair of athletic-type runners or similar), with the addition of orthotics if usually worn. The course was a 20-metre-long circuit on a flat vinyl surfaced corridor in a hospital outpatient clinic. In Sydney, participants were assessed barefoot unless they required ankle foot orthoses (AFOs), in which case they wore AFOs with their usual footwear. The course was a 25-metre-long circuit on a flat carpeted surface in a hospital outpatient clinic. Six-minute walk distance (6MWD) was normalised to height to account for differences in height across the large age range of participants (26).

### Walk-12

Participants with CMT completed a Walk-12 questionnaire (27), with participants aged 8 years and younger assisted by a parent or adult. The Walk-12 is a validated self-reported questionnaire used to measure the child’s experience and perception of the limitations of their disease on their gait and gait-related activities (27). Briefly, 12 statements relating to gait and gait-related activities were answered on a 5-point Likert scale from zero (0) “my disease does not affect this” to five (5) “I am very limited by my disease”. The raw score out of 60 was transformed to provide a composite score; a higher score indicating greater impact of disease on walking ability (28).

### Data management

Data from the clinical assessments were recorded on paper case report forms (CRF) and subsequently entered in an electronic data management system (REDCap™), hosted by the Murdoch Children’s Research Institute (29). Data from the two questionnaires (PAQ-C and Walk-12) were either completed on a paper CRF or entered directly into REDCap™ by the participants during the study visit. Paper CRFs from Sydney were scanned and emailed to Melbourne for data entry.

### Sample size calculation and statistical analysis

Assuming normality of the PAQ-C data, a sample size of 50 in each group with α_1_ = 0.05 and a moderate effect size (d = 0.50) allowed for >80% power to detect differences between the groups. When considering the power of the correlation co-efficient analysis a sample of 50 with r = 0.5, α_1_ = 0.05 allowed for >90% power to detect associations between the CMTPedS, Walk-12 and PAQ-C.

Normality of data distribution was visually examined and statistically tested for all variables with Shapiro-Wilk test. Appropriate descriptive statistics were used to characterise the data. Based on the type and distribution of the data, parametric (*t*-test) or non-parametric (Wilcoxon signed-ranks test) tests were used to assess for differences between the CMT and TD groups. Effect size was calculated with Cohen’s *d*. For the 6MWD analysis, only participants who wore footwear (with or without orthoses) were included, as footwear is known to positively impact gait speed, step length and therefore 6MWD, compared to barefoot gait (1, 30). Spearman’s correlations were used to determine associations between physical activity (PAQ-C), and disease severity (CMTPedS), ambulatory capacity (normalised 6MWD) and perceived walking ability (Walk-12). Stata v15 was used for data analysis.

## Results

### Participants

One hundred children, 50 with CMT and 50 age- and gender-matched TD controls, participated in this study with a mean age of 12.5 years (range 4-18 years old). CMT subtypes included CMT1A (n=37), CMTX (n=6), CMT2 (n=2), CMT1B (n=1), CMT4A (n=1), CMTDIB (n=1) and unknown/no genetic diagnosis (n=2). Participant characteristics are outlined in Table 1. There were no statistical differences in height and weight between the groups. The CMT group had a higher BMI compared to the TD group (*p* = 0.003) but remained within a healthy range of 16-21 kg/m^2^ for the mean age of the study groups (31). The participants with CMT were moderately affected by their disease, had weaker ankle dorsiflexor muscles, poorer balance and reduced lower limb muscle power as assessed with a standing long jump (all *p* < 0.001) compared to their TD peers.

**Table 1.** Participants’ characteristics and clinical measures (CMT and TD.

|  | **CMT**<br>**Mean (SD) [95% CI]** | **TD**<br>**Mean (SD) [95% CI]** | **Mean difference** | **Effect size**<br>*d* | *p* value |
| --- | --- | --- | --- | --- | --- |
| **Gender male : female** | 24:26 | 24:26 | - | - | - |
| **Age**<br>**(years)** | 12.5 (3.9)<br>[11.4, 13.6] | 12.5 (3.9)<br>[11.4, 13.6] | - | - | - |
| **Height**<br>**(m)** | 1.51 (0.20)<br>[1.46, 1.57] | 1.51 (0.20)<br>[1.46, 1.57] | - | - | 0.9 |
| **Weight**<br>**(kg)** | 49.1 (21.8)<br>[43.0, 55.3] | 44.5 (17.2)<br>[39.6, 49.4] | 4.7 (17.9)<br>[0.4, -9.8] | - | 0.07 |
| **BMI**<br>**(kg/m<sup>2</sup>) †** | 20.5 (5.9)<br>[18.8, 22.2] | 18.6 (3.5)<br>[17.5, 19.6] | 2.0 (5.7)<br>[0.4, 3.6] | - | 0.003 |
| **Max dfl range**<br>**(degrees)** | 25.0 (6.2)<br>[23.1, 26.9] | 36.7 (7.6)<br>[34.4, 39.1] | 11.7 (1.5)<br>[8.7, 14.7] | 1.7 | < 0.001 |
| **FPI - left**<br>**median (IQR)**<br>**(-12 to +12) †** | 0<br>(-2 to 6) | 5<br>(3 to 6) | 5 | - | < 0.001 |
| **FPI - right**<br>**median (IQR)**<br>**(-12 to +12) †** | 1<br>(-2 to 6) | 5<br>(2 to 6) | 4 | - | 0.01 |
| **Dflr strength**<br>**(Newtons) <sup>a</sup>** | 47.3 (27.4)<br>[39.5, 55.1] | 90.1 (25.6)<br>[82.8, 97.4] | 42.8 (37.6)<br>[39.5, 55.1] | 1.6 | < 0.001 |
| **Plflr strength**<br>**(Newtons) <sup>a</sup>** | 146.0 (76.5)<br>[124.2, 167.7] | 153.2 (43.7)<br>[140.8, 165.6] | 7.2 (82.2)<br>[-16.1, 30.6] | 0.12 | 0.5 |
| **CMTPedS**<br>**( /44)** | 20 (8)<br>[18, 23] | - | - | - | - |
| **Balance BOT**<br>**( /37) <sup>a</sup>** | 19 (9)<br>[17, 22] | 32 (3)<br>[31, 33] | 12.1 (9.2)<br>[9.5, 14.7] | 1.8 | < 0.001 |
| **Standing long jump**<br>**(m) <sup>a</sup>** | 0.69 (0.38)<br>[0.58, 0.80] | 1.42 (0.32)<br>[1.33, 1.51] | 0.73 (0.50)<br>[0.59, 0.87] | 2.0 | < 0.001 |
| **6MWD (footwear)**<br>**(m) <sup>a b</sup>** | 485.1(160.9)<br>[427.1, 543.1] | 639.8 (83.1)<br>[609.9, 669.8] | 154.8 (170.4)<br>[93.3, 216.2] | 1.2 | < 0.001 |
| **N6MWD (footwear) <sup>b c</sup>** | 338.4 (114.6) | 440.9 (54.0) | 102.5 (95.4) | 1.1 | < 0.001 |
|  | [297.1, 379.8] | [421.4, 460.4] | [68.1, 136.8] |  |  |

† Wilcoxon signed-ranks test; <sup>a</sup> selected lower limb items from the CMTPedS completed by TD participants; <sup>b</sup> 6MWT in footwear n = 32; <sup>c</sup> normalised to height. Abbreviations: 6MWD: six minute walk distance; BMI: body mass index; BOT: Bruininks-Oseretsky test of motor proficiency; CMT: Charcot-Marie-Tooth disease; CMTPedS: CMT pediatric scale; Dflr: dorsiflexor; ES: effect size; FPI: Foot posture index; IQR: interquartile rank; Max dfl range: maximum dorsiflexion range; N6MWD: Normalised 6MWD to height; Plflr: plantar flexor; TD: typically developing

### Measure of physical activity

Participants with CMT were less physically active than their TD peers as reported on the PAQ-C (*p* < 0.001; Table 2). The CMT reported group estimate of MVPA was 40 minutes per day (estimated MVPA 283.6 mins per week), whereas the TD reported group estimate was 45 minutes per day (estimated MVPA 318.0 mins per week) (*p* < 0.001). One participant with CMT did not complete the PAQ-C questionnaire; their matched TD control was removed from the analysis.

**Table 2.** Physical activity score (PAQ-C) and estimated weekly moderate to vigorous physical activity for the CMT and TD groups.

|  | **CMT n = 49** | **TD n = 49** | ***p* value** |
| --- | --- | --- | --- |
| **PAQ-C † (/5)** | 2.2 | 2.8 | < 0.001 |
| **Median (IQR)** | (1.5–2.7) | (2.3–3.3) |  |
| **Estimated MVPA (mins/week)** | 283.6 (211.6) | 318.0 (202.5) | < 0.001 |
| **Mean (SD) [95% CI]** | [222.8, 344.3] | [257.2, 374.4] |  |

† Wilcoxon signed-ranks test. Abbreviations: CMT: Charcot-Marie-Tooth disease; TD: typically developing; PAQ-C: physical activity questionnaire child; MVPA: moderate to vigorous physical activity; IQR: interquartile range; SD: standard deviation; CI: confidence interval

### Measures of functional ambulation

Thirty-two participants with CMT completed the 6MWT in footwear and 18 walked barefoot due to site dependent protocol differences. Seven participants with CMT wore AFOs and two wore customised foot orthoses. No TD participant wore orthoses. Analysis of the 6MWD included only the subset of 32 participants with CMT who wore footwear and their TD matched controls.

Participants with CMT walked significantly shorter distances in six minutes compared to their TD peers with a large effect size (mean difference 154.8 m, *p* < 0.001, *d* = 1.2 (95% CI [93.3, 216.2]); Table 1). When 6MWD was normalised to the participant’s height to account for the influence on step length, the significant difference remained, as did the large effect size (mean difference 102.5, *p* < 0.001, *d* = 1.1 (95% CI [68.1, 136.8]); Table 1).

Participants with CMT reported that their disease generally had a mild effect on their gait and gait-related activities with a transformed Walk-12 mean score of 24.7% (mean 24.7 (SD 19) 95% CI [19.3, 30.2]). Closer inspection of the individual questions revealed that except for the use of gait aids, 50-75% of the participants felt that their disease affected their gait and gait-related activities at least “a little bit” (Fig 1). Nearly 30% or more of participants reported moderate or greater effects on their ability to run (43%), ascend or descend stairs (41%), how far they could walk (41%), increased concentration required when walking (39%), the smoothness or co-ordination of their walking (33%), increased effort to walk (31%) and how fast they could walk (31%) (Fig 1). One participant with CMT did not complete the Walk-12 questionnaire.

**Fig 1.**
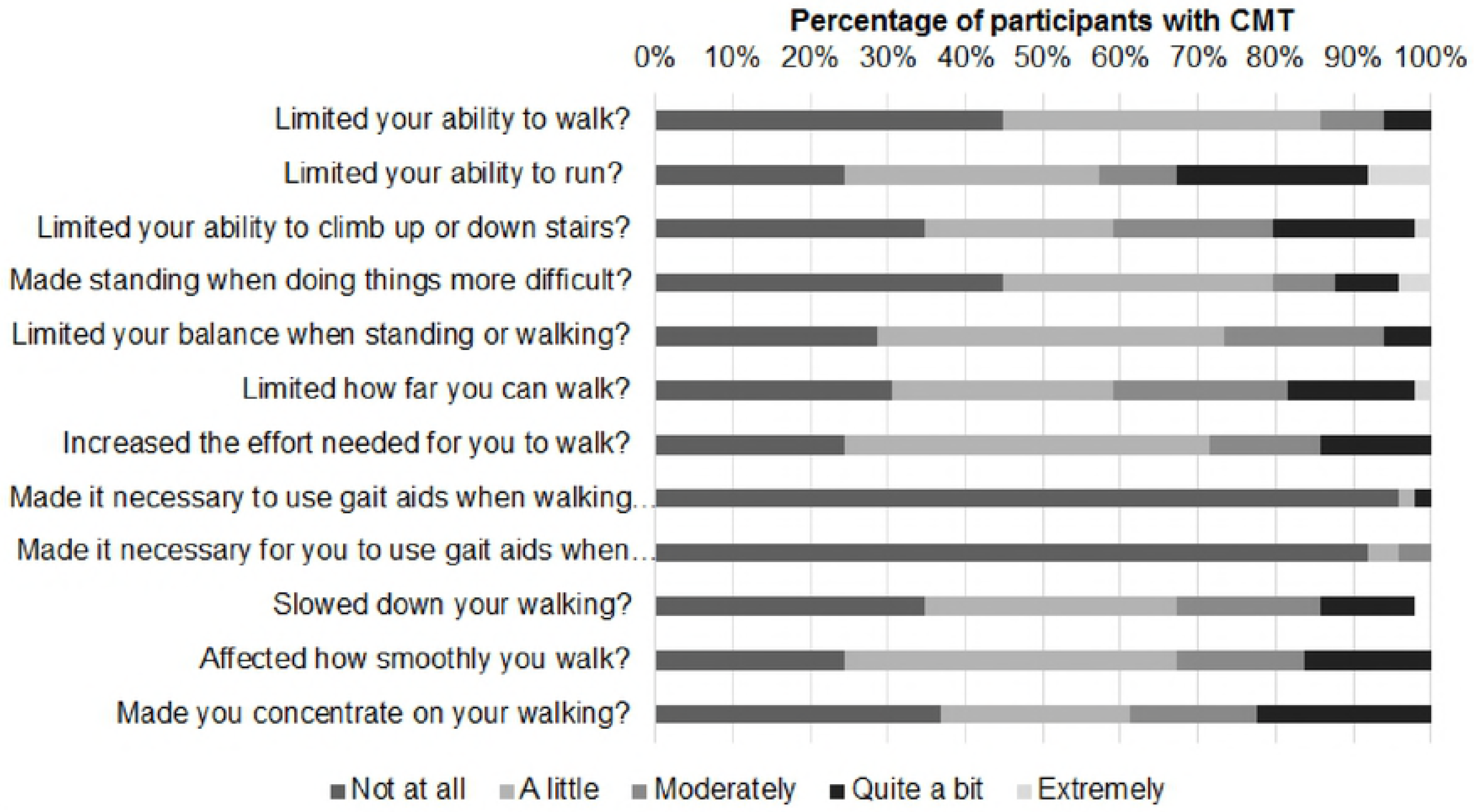
Percentage of participants with CMT (n=49) responses to the question “How much has your disease…”.

### Associations between physical activity, disease severity, functional ambulation and perception of disease effect on gait and gait-related activities

Spearman’s correlation analyses between physical activity (PAQ-C), disease severity (CMTPedS), functional ambulatory capacity (normalised 6MWD – footwear) and perception of disease effect on gait and gait-related activities (Walk-12) in 31 participants with CMT are presented in Table 3. The strongest association was between physical activity and normalised 6MWD with a moderate to good association between the two (ρ = 0.71, *p* < 0.001) and an inverse moderate association between physical activity and disease severity (ρ = -0.52, *p* < 0.001). The moderate to good association between normalised 6MWD and CMTPedS was expected given that the 6MWD is a scored item in the CMTPedS (ρ = -0.78. *p* < 0.001).

**Table 3.**
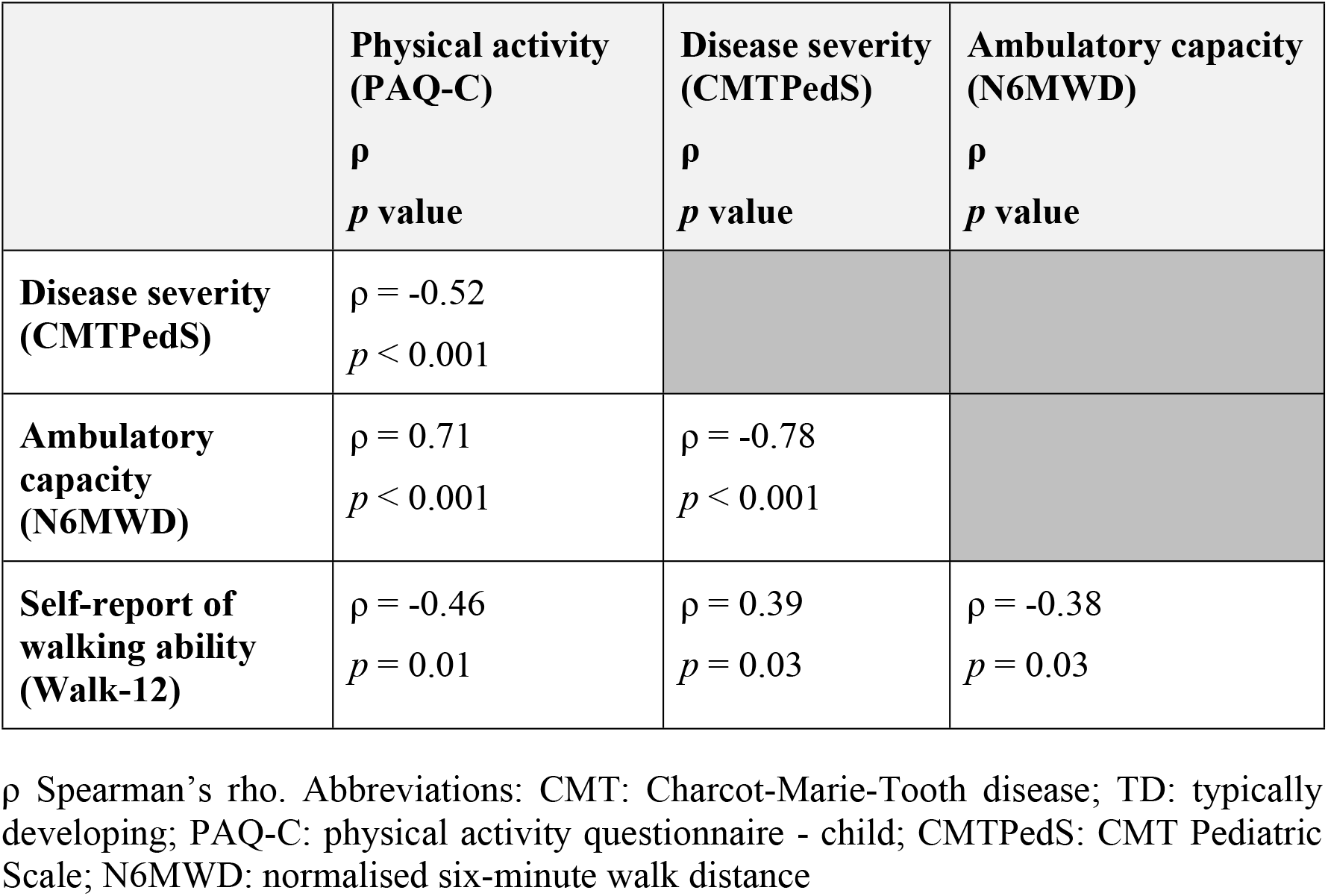
Correlations between physical activity (PAQ-C), disease severity (CMTPedS), functional ambulatory capacity (normalised 6MWD) and self-reported effect of disease on gait and gait-related activities (Walk-12) in 31 participants with CMT.

| | **Physical activity**<br>**(PAQ-C)**<br>$\rho$<br>$p$ value | **Disease severity**<br>**(CMTPedS)**<br>$\rho$<br>$p$ value | **Ambulatory capacity**<br>**(N6MWD)**<br>$\rho$<br>$p$ value |
| --- | --- | --- | --- |
| **Disease severity**<br>**(CMTPedS)** | $\rho = -0.52$<br>$p < 0.001$ | | |
| **Ambulatory**<br>**capacity**<br>**(N6MWD)** | $\rho = 0.71$<br>$p < 0.001$ | $\rho = -0.78$<br>$p < 0.001$ | |
| **Self-report of**<br>**walking ability**<br>**(Walk-12)** | $\rho = -0.46$<br>$p = 0.01$ | $\rho = 0.39$<br>$p = 0.03$ | $\rho = -0.38$<br>$p = 0.03$ |

$\rho$ Spearman's rho. Abbreviations: CMT: Charcot-Marie-Tooth disease; TD: typically developing; PAQ-C: physical activity questionnaire - child; CMTPedS: CMT Pediatric Scale; N6MWD: normalised six-minute walk distance

## Discussion

The findings of this study across two Australian paediatric neuromuscular centres suggest that children with CMT report being less physically active than their TD peers. Of note, nearly 75% of the children in this study were diagnosed with CMT1A, yet despite the high representation of this generally milder CMT sub-type, reported levels of physical activity and functional ambulation were considerably reduced (32). Lower physical activity levels were associated with greater disability and reduced functional ambulation, both in terms of ambulatory capacity in a 6MWT and perceived disease-related limitations to gait and gait-related activities.

In this study we found that children with CMT are less active. It is therefore important in the context of a degenerative disease with increasing physical impairment to monitor physical activity levels in children with CMT. Reduced physical activity is known to increase health-related illnesses in the general population and is likely to compound the effect of disease. Maintaining general health and wellness is important for participation in education, paid employment and general activities of daily life. Understanding the importance of engaging and participating in physical activity whilst living with a physical disability is an important health message that health clinicians working with children with CMT need to promote. It is important that the link between greater physical activity and better function is understood. The evidence from this study suggests that children with CMT who are more physically active, have milder disease and greater capacity to undertake gait-related activities. Conversely, children with greater disease severity are less able to engage in physical activity leading to increased sedentary activity time. This may have a downward spiralling effect further reducing physical activity with greater impact on disease and ambulatory function.

Children who reported being less physically active also demonstrated reduced ambulatory capacity. This has implications for community mobility where distance requirements in community settings can range up to 700 metres (33). Children who are more disabled are further limited in their ability to access community activities that require longer distances to be traversed. The children with CMT who were less active had greater disease severity and reported greater limitations to their gait-related activities, similar to the findings of a small study of children with mixed neuromuscular diagnoses (34). Prior research suggests this may be a lifelong concern, as adults with CMT also report limitations to physical activity (8, 9). The current study expands on the studies in adults to establish that physical activity is limited from childhood in CMT, similar to other paediatric disorders that present with physical impairments such as cerebral palsy, muscular dystrophy and spina bifida (7).

It is uncertain what factors may facilitate or present as barriers to physical activity in children with CMT. Some evidence from children with other physical disabilities indicates that they may self-limit physical activity behaviours to avoid social embarrassment in sporting and social situations (35). Similarly, it is likely that the risk of social embarrassment may also be a potential barrier to physical activity in children with CMT. A previous study found that children with CMT are at a higher risk of falling (36). The current study identified that children with CMT have impaired balance; over 60% considered that their walking was uncoordinated or less smooth and that they needed to concentrate on their walking. Children with CMT may self-limit physical activity to reduce and prevent falling or near-falls, similar to reports in young adults with cerebral palsy (37). Further barriers may include difficulty accessing activities that are inclusive and provide opportunities to be physically active in the local community (38). Sourcing disability-trained and supportive trainers or coaches to adapt and modify activities for children with a physical disability is an ongoing problem in the Australian context (38).

There were several strengths to this novel study of physical activity in children with CMT. A well-described CMT cohort included a broad age and developmental range spanning childhood and adolescence and comprised CMT sub-types representative of the clinical populations from two specialist paediatric neuromuscular centres. Disease severity was well-characterised with the CMTPedS, a meaningful and validated measure of disability in paediatric CMT. Generalisability to other CMT populations is likely, given that disease severity in this study was similar to reported levels in a large international cohort of 520 children (32). The age- and gender-matching to TD controls strengthened the study and placed the findings within the context of typical development.

Utilisation of the calibration method of estimating MVPA suggested by Saint-Maurice et al. (2014) provided a meaningful quantification of physical activity however it was limited to a group estimate only. Further, limitations of the self-report PAQ-C is that it relies on recall memory and the child’s perception of what constitutes physical activity. Self-report recall physical activity questionnaires are therefore less accurate compared to objective measures (12). The use of an activity monitor or accelerometer would have provided a more robust measure of MVPA cut-points and enabled further comparison to National Physical Activity Guidelines (39). Additionally, the PAQ-C did not measure sedentary time. The clinical outcome measures were collected per local protocols and limited the 6MWT analysis. Differences in 6MWT circuit length were unlikely to affect group 6MWD, as the typically developing cohort were tested on the shorter circuit. Therefore, any error due to circuit length and a greater number of turns was likely to have underestimated the group difference (40).

Children with CMT are at risk of greater health problems and impairment due not only to their disease but also disuse associated with reduced physical activity. Objective quantifiable assessment of physical activity in paediatric CMT, and investigation of potential barriers to physical activity including fear of falling, are areas that require further enquiry. Further investigation of enablers of physical activity is suggested, including the effects of exercise and increased physical activity on ambulatory function in children with CMT. Strength training improves activities of daily living in adults with CMT (41), and is safe in children with CMT (42). In TD children, increasing physical activity from one 60-minute physical education class a week to five 40-minute classes a week improved muscle strength (43). Given these findings, the effectiveness of age-appropriate regular exercise programs, utilising motivational behaviour change coaching and focussing on strength training and whole-body activity for children with CMT, requires further development and empirical investigation. Additionally, facilitating access to and participation in community-based recreational and sporting activities, including specialist training to enable coaches to adapt programs to include children with physical disabilities, is required.

## Conclusion

Physical activity and functional ambulation are adversely and significantly affected, and are associated with greater disability, in children with CMT. Healthcare clinicians, researchers and funding agencies ought to engage with and promote opportunities for children with CMT to be more physically active, be it through participation in structured, evidence-based exercise and training programs, or community-based recreational sporting programs. Further research is required to determine whether facilitating greater physical activity may slow disease progression and improve physical function in the everyday lives of children and adolescents with CMT.

### List of abbreviations

6MWT: six-minute walk test
6MWD: six-minute walk distance
AFOs: ankle foot orthoses
BMI: body mass index
CMT: Charcot-Marie-Tooth disease
CMTPedS: CMT Pediatric Scale
CRF: case report form
MVPA: moderate to vigorous physical activity
N6MWD: normalised six-minute walk distance
PAQ-A/-C: physical activity questionnaire – adolescent/child
TD: typically developing

## Acknowledgements

The authors would like to acknowledge Dr Kayla Cornett and Katy de Valle for assistance with data collection in Sydney and Melbourne respectively. Further thanks to the staff and patients of the Neuromuscular Clinic, The Royal Children’s Hospital and the Peripheral Neuropathy Management Clinic, Children’s Hospital, Westmead.

**Authors’ contributions**
RAK, JLM, KC, KLP and JB were involved in the study conceptualization and design. MMR and JB provided study resources including clinic patients. RAK, KC and KR were responsible for recruitment and assessment of participants. RAK was responsible for data analysis and original draft preparation. All authors were responsible for review and editing of the manuscript.

